# Understanding the role of membrane lipids in mechanism of antimicrobial photodynamic therapy in Escherichia coli

**DOI:** 10.64898/2026.01.26.701284

**Authors:** Marta Piksa, Mariusz A. Bromke, Carlos M. Marques, Sigolène Lecuyer, Patricia Daira, Baptiste Fourmaux, Ifor D. W. Samuel, Katarzyna Matczyszyn, Krzysztof J. Pawlik

**Affiliations:** Hirszfeld Institute of Immunology and Experimental Therapy, Poland; Wroclaw Medical University, Department of Biochemistry and Immunochemistry, Poland; Ecole Normale Supérieure de Lyon, France; Institut National des Sciences Appliquées de Lyon, France; University of St Andrews, Scotland; Wrocław University of Science and Technology, Poland

## Abstract

Antimicrobial photodynamic therapy (aPDT) is a promising alternative to antibiotics, yet the molecular factors determining bacterial susceptibility remain unclear.

This study investigates the critical role of bacterial lipids, particularly cardiolipin (CL) in the aPDT response of *Escherichia coli* using methylene blue as a photosensitizer.

Through genetic deletion (ΔclsABC) and chemical modification via mannitol supplementation, we demonstrate that reduced CL levels significantly enhance bacterial sensitivity, leading to an additional reduction in viability exceeding 3 log10. Quantitative lipidomics (MS,GC) confirmed substantial CL depletion and altered fatty acid saturation. Interestingly, while live CL-deficient cells were more vulnerable, biomimetic giant unilamellar vesicles (GUVs) with higher CL content showed greater susceptibility to photo-oxidation. These findings suggest that CL-rich microdomains in living bacteria act as functional scaffolds for stress-defense systems rather than mere targets for oxidative damage. Modulating membrane lipid composition thus represents a novel strategy to potentiate aPDT efficacy.

## Introduction

Photodynamic therapy (PDT) has a rich history, spanning over a century, with its application in treating pathogens gaining momentum in recent decades. The term *photodynamic action* was introduced in the beginning of 20th century to describe the light-activated process involving photosensitizers, light, and molecular oxygen.^1,2^ This interaction results in the generation of reactive oxygen species (ROS), leading to oxidative stress, characterized by an imbalance between the ROS production and antioxidant defences. Oxidative stress plays a pivotal role in cellular damage by oxidizing key biomolecules such as lipids, proteins, and nucleic acids.^3,4^

Antimicrobial photodynamic therapy (aPDT), as a tool for fighting pathogens, is being extensively explored primarily due to its potential to address challenges posed by antibiotic resistance, the increasing complexity of infections, and the need for more targeted antimicrobial therapies.^5,6^ One of the major advantage of aPDT lies in its spatial precision. When combined with light of specific wavelengths, aPDT can target only the infected areas without disturbing surrounding healthy tissue.^1,7^ This precision reduces side effects, making aPDT particularly attractive for treating infections in sensitive areas, such as in dentistry, wound care, or dermatological infections.^8–11^ In addition, photodynamic therapy exhibits broad-spectrum antimicrobial activity, effectively targeting bacteria, fungi, and viruses.^12^ This versatility opens up opportunities for treating various types of infections. Furthermore, aPDT has proven ability to disrupt microbial biofilms, a common contributor to persistent and chronic infections.^13–15^

As we have shown previously, the effectiveness of aPDT varies between pathogens, influenced by factors such as the initial target amount, irradiation time, photosensitizer concentration, the physiological state of the microbes, and their structural and biochemical characteristics.^6,12,16^ A primary hypothesis attributes these variations in susceptibility to fundamental differences in bacterial cell wall structure. Gram-positive bacteria (G+) are usually more sensitive to aPDT than Gram-negative bacteria (G−), even though G+ have a thicker peptidoglycan layer. In contrast G− bacteria possess an additional outer membrane rich in negatively charged lipopolysaccharides, which can hinder aPDT effectiveness. Nonetheless, even within the same genus, variability in response is broadly observed^6,16^ The precise molecular and biochemical determinants underlying this phenomenon remain largely unclear. Among all biomolecules susceptible to oxidative damage during PDT, lipids are unique in undergoing a chain reaction mechanism of oxidation. Within the diverse group of lipids constituting the cell membrane, one in particular stands out due to its susceptibility to oxidation - cardiolipin (CL), also known as bisphosphatidylglycerol. This specific anionic phospholipid, present not only in the mitochondrial membranes of eukaryotic cells but also in the membranes of several bacterial species, is considered to play a crucial role in the context of PDT. ^17–20^ In eukaryotic cells, cardiolipin oxidation leads to mitochondrial dysfunction and triggers apoptosis - a mechanism well-characterized in cancer photodynamic therapy.^21–23^ In *E. coli*, cardiolipin is predominantly localized at the pole regions of the inner membrane and constitutes approximately 5-10% of the total phospholipid content.^18,24^ It has emerged as a potential vulnerability factor due to its involvement in maintaining membrane curvature, optimal membrane protein translocation and insertion, controlling the osmotic stress response, DNA replication, respiratory chain and enzymatic activity of bacterial cytochromes.^25–29^ Impairment of cardiolipin production in the bacterial cell membrane can be achieved either genetically or chemically. ^30,31^ Deletion of cardiolipin synthase genes *clsA, clsB* and *clsC* results in significant decrease of cardiolipin level in *Escherichia coli*. Notably, supplementing bacterial cultures with high concentrations of mannitol leads to great changes in cardiolipin profile by induction of the biosynthesis of novel lipids. In particular the glycerol backbone may be substituted with mannitol leading to phosphatidylmannitol and diphosphatidylmannitol. Such modification results in lipid remodelling potentially affecting membrane dynamics and susceptibility of bacteria to aPDT. Understanding the role of lipids, particularly cardiolipin, in antimicrobial photodynamic therapy opens new possibilities for enhancing treatment specificity and efficacy, especially against Gram-negative bacteria such as *Escherichia coli*, which are typically more resistant to aPDT. Elucidating the involvement of cardiolipin also represents a significant step toward a deeper understanding of the fundamental mechanisms underlying aPDT in bacterial cells.

The aim of the study is to provide a comprehensive insight into the essential role of cardiolipin in photodynamic antibacterial therapy, using *E. coli* bacteria as a model organism, and to evaluate the impact of changes in lipid composition on the susceptibility of microorganisms to aPDT. Methylene blue (MB) was selected as the photosensitiser for the research due to its clinical approval, water solubility, minimal toxicity in the dark, and widespread availability. To accurately unravel the role of cardiolipin and associated lipid profile changes in the context of aPDT, the study will integrate measurements of bacterial viability and cell membrane integrity with detailed quantitative lipidomic analysis and identification of cardiolipin microdomains in live bacterial cells. This comprehensive, multi-faceted approach is essential for investigating the direct relationship between specific biochemical changes in lipid membranes and the resulting bactericidal efficacy of aPDT, which will help to explain the observed changes in microbial sensitivity.

## Results

### APDT on *Escherichia coli* with impaired cardiolipin production

The efficiency of aPDT was evaluated on planktonic bacterial cultures with a chemically modified membrane lipid composition, by growing them in the presence of 0.6 M D-mannitol, and *E. coli ΔclsABC* mutant. The final concentration of photosensitizer methylene blue (MB) in the bacterial suspension was 15 μM. Results of the experiments demonstrate a significant variation in aPDT efficacy depending on the membrane lipid composition. Bacteria subjected to aPDT under standard growth conditions (no chemical treatment or genetic modification) exhibited a reduction in viability of 1.75 ± 0.49 log10 (mean ±standard deviation). In contrast, culture modified with 0.6 M D-mannitol supplementation during aPDT led to an additional reduction of bacteria by more than 3 orders of magnitude, reaching 5.32 ± 1.14 log10. A similar enhancement to that in chemically modified culture was observed in the *E. coli ΔclsABC* mutant lacking cardiolipin due to the triple deletion of *clsA, clsB* and *clsC* genes responsible for cardiolipin biosynthesis. This targeted mutation resulted in a significantly higher bacterial reduction by aPDT equal to 3.44 ±0. 34 log10 compared to the control. Furthermore, the mutant strain exhibited similarly increased sensitivity to methylene blue, with a reduction of 2.12 ± 0.22 log10 (Fig.1)

**Fig 1.**
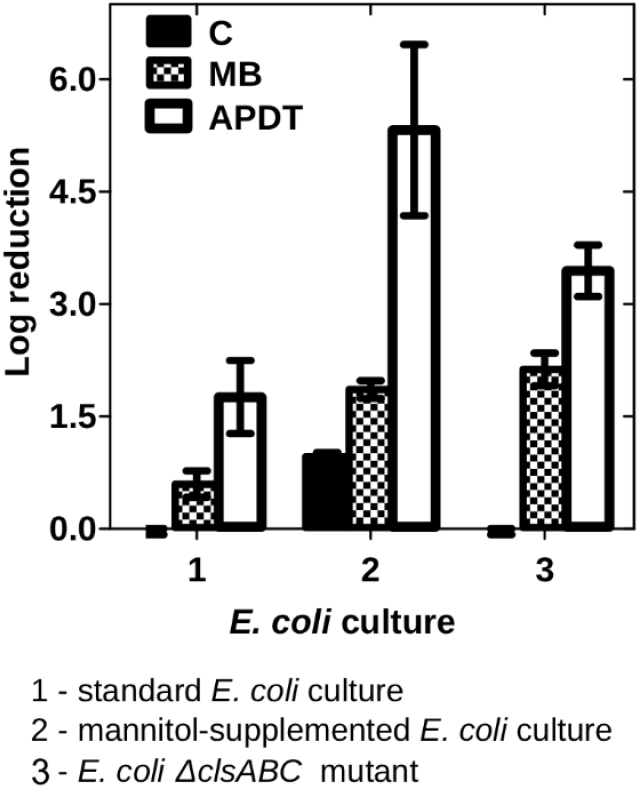
Effects of antimicrobial photodynamic therapy (aPDT) and methylene blue (MB) on *Escherichia coli*. Three experimental groups were analysed: 1-standard *E. coli* culture – untreated *E. coli* grown under standard conditions, 2 - chemically modified by 0.6 M D mannitol presence *E. coli* culture, 3 - *E. coli ΔclsABC* mutant – *E. coli* strain lacking cardiolipin due to the deletion of the *clsABC* genes. The results are presented as follows: empty bars represent aPDT treatment (exposure to methylene blue (MB) and light), patterned bars represent MB treatment (methylene blue exposure, no light), black bars represent the control (C) (no methylene blue or light exposure). Each bar shows the mean of three biological replicates, each with three technical repeats (n=9). Error bars indicate the standard deviation. Statistical analysis (t-test) and data visualization were performed using GraphPad Prism.

The concentration of D-mannitol used had a noticeable effect on bacterial growth in the absence of irradiation or MB, causing a moderate reduction of 0.96 ± 0.05 log10. Additionally, the presence of D-mannitol enhanced bacterial susceptibility to MB alone, resulting in a reduction of 1.86 ± 0.11 log10, compared to 0.59 ± 0.17 log10 in untreated cultures. To exclude the influence of osmotic stress, control aPDT experiments were performed using 0.33M NaCl - a concentration showing identical growth inhibition to 0.6 M mannitol. Under these conditions no change in aPDT-induced reduction and no increase in methylene blue influence was observed (data not-shown).

### The electrokinetic properties of bacterial cell surface

Alterations in the bacterial lipid profile are examined through measurements of electrokinetic membrane properties, which provide insights into surface charge and membrane integrity, parameters that critically determine bacterial permeability, stability, viability, and interactions with antimicrobial agents or other bioactive molecules. The measurements of the bacterial surface charge exhibited significant differences in electrokinetic membrane properties depending on the culture conditions. The lipid composition change by D-mannitol presence in the medium notably altered electrokinetic properties of *E. coli*, in particular causing a positive shift in the Zeta Potential (ZP) and thereby decreasing the magnitude of the net negative surface charge. Electrophoretic mobility increased 1,9-fold in the mannitol-supplemented culture compared to the standard condition (Fig. 2b), with corresponding rise in ZP values (Fig. 2c). In contrast, performed modification of lipid composition does not significantly influence the culture conductivity. Both cultures of *E*.*coli* retained the ability to conduct electric current at nearly the same level (Fig. 2a). No substantial change in the electrical conductivity of the culture medium with or without D-mannitol is observed, indicating stable ion concentrations in both conditions and confirming that the ZP change is due to an alteration of the cell modification, rather than a change in the surrounding medium.

**Fig 2.**
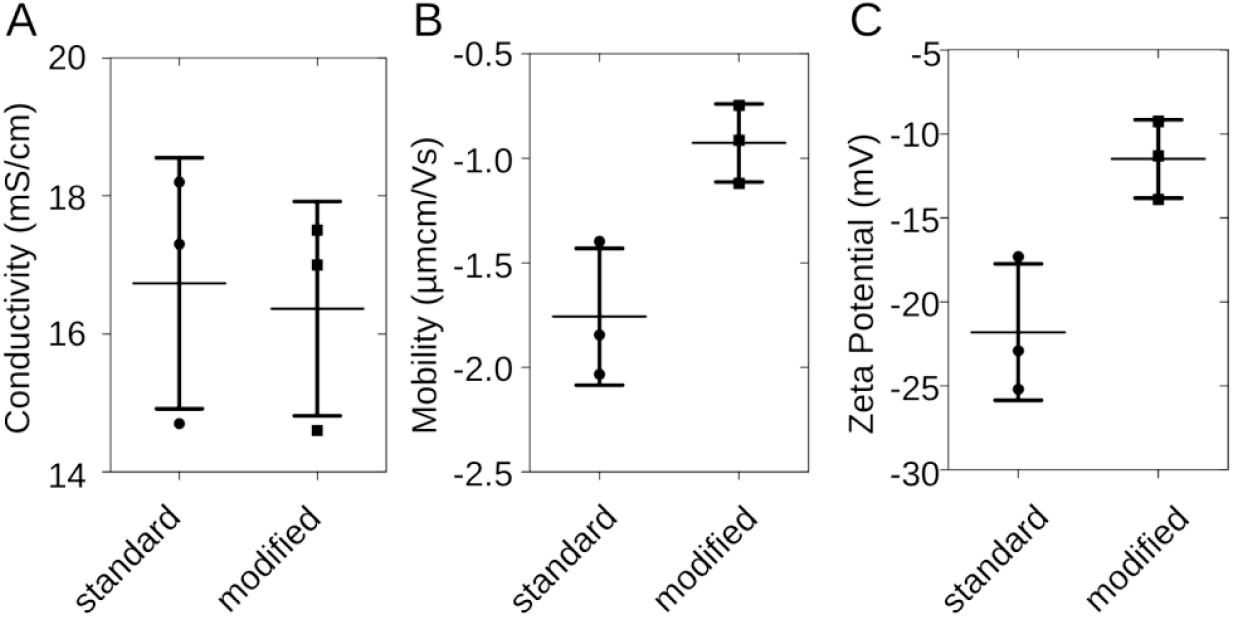
Effect of on electrokinetic properties of *E. coli*. The graph presents data on conductivity (A), mobility (B), and zeta potential (C) in *E. coli*: standard culture and chemically modified culture. Each horizontal line represents the average value from three independent biological replicates, with each replicate measured three times (n=9). The whiskers indicate the standard deviation. Statistical analysis and data visualization were conducted using GraphPad Prism.

### Photodynamic response of biomimetic bacterial membrane model

Lipid extracts from *E. coli* grown under two distinct conditions: (1) standard - untreated *E. coli*, and (2) chemically modified - *E. coli* grown in the presence of 0.6 M D-mannitol, were used to construct biomimetic membrane models in the form of giant unilamellar vesicles (GUVs). Lipids were extracted independently from each culture and used for GUV preparation, followed by morphological and photochemical evaluation using phase-contrast microscopy. The vesicles response to photosensitisation (photosensitizer and light) was analysed, revealing no significant differences in their morphology or size (Fig. 3A, D), the photochemical response exhibited a marked alteration by modifying *E. coli* with D-mannitol. Under 15 μM MB and red-light exposure, GUVs assembled from standard *E. coli* lipid extracts (n=40) showed a sharp decrease in phase contrast between the vesicle lumen (100 mM sucrose solution) and the surrounding medium (90 mM glucose solution) (Fig. 3B, C). The contrast variation with irradiation time is characterized by an initial plateau where the contrast does not change, followed after a certain lag time (11±7 seconds measured on 40 vesicles) by a sudden decrease over a transition time 22±15 seconds. This change in phase contrast is indicative of alterations in the vesicle structure, such as increased permeability, leakage of internal contents, or changes in internal osmotic balance, and suggests a reduction in the internal solute concentration, leading to the equalization of the refractive index between the lumen and the medium. The observed delay in the onset of significant phase contrast decrease, followed by a sustained reduction, highlights the dynamic nature of membrane damage (Fig. 3C). The individual traces show heterogeneity in the onset and rate of phase contrast change among vesicles, which may reflect variations in vesicle size, photosensitizer incorporation, or local light intensity. In contrast, GUVs derived from lipid extracts of modified cultures maintained their structural integrity and optical contrast, even after 18 minutes of illumination (Fig. 3A, C, D), suggesting increased resistance to aPDT condition.

**Fig 3.**
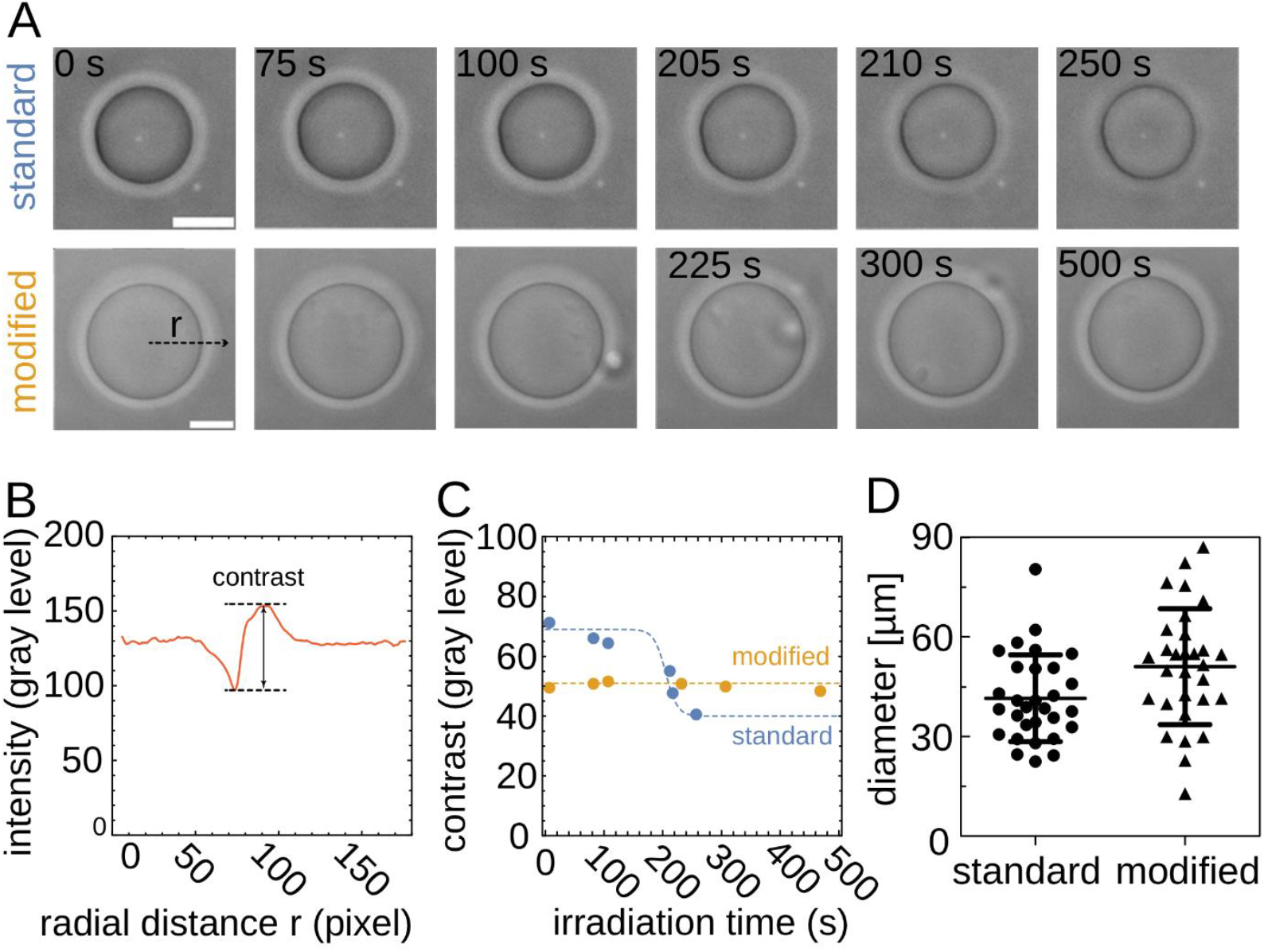
Sensitivity to photo-induced oxidation of GUVs assembled from bacterial extracts. (A) Phase contrast images of GUVs in 15 μM MB under constant irradiation (scale bars: 20 μm). (B) A measure of the contrast is provided by the maximum difference between gray levels extracted across the radial profile (C) Contrast evolution for both types of vesicles, showing that GUVs assembled from lipid extracts from chemically modified bacteria are less prone to oxidation. (D) Mean diameters for both types of GUVs (n = 40 each group).

### Cardiolipin visualisation by fluorescence microscopy

Incubation with 10-nonyl bromide acridine orange (NAO, 200 nM) allowed the visualization of cardiolipin-rich microdomains in *Escherichia coli* (Fig. 4). The final effect was strongly dependent on both, the culture conditions and the immobilisation method used. In mannitol-supplemented culture, no NAO-derived fluorescence signal was detected, regardless of the immobilization matrix applied (Fig. 4A3, A.4). This observation confirms the absence of cardiolipin in the modified membrane. Conversely, cells from standard cultures exhibited extensive green fluorescence associated with NAO binding (Fig. 4A1, A2, B). Further analysis using poly-L-lysine-mediated immobilization allowed for the visualization of specific regions within *E. coli* cells, where NAO staining and fluorescence was strongly associated with cardiolipin microdomains, particularly at the cell poles (Fig. 4B).

**Fig 4.**
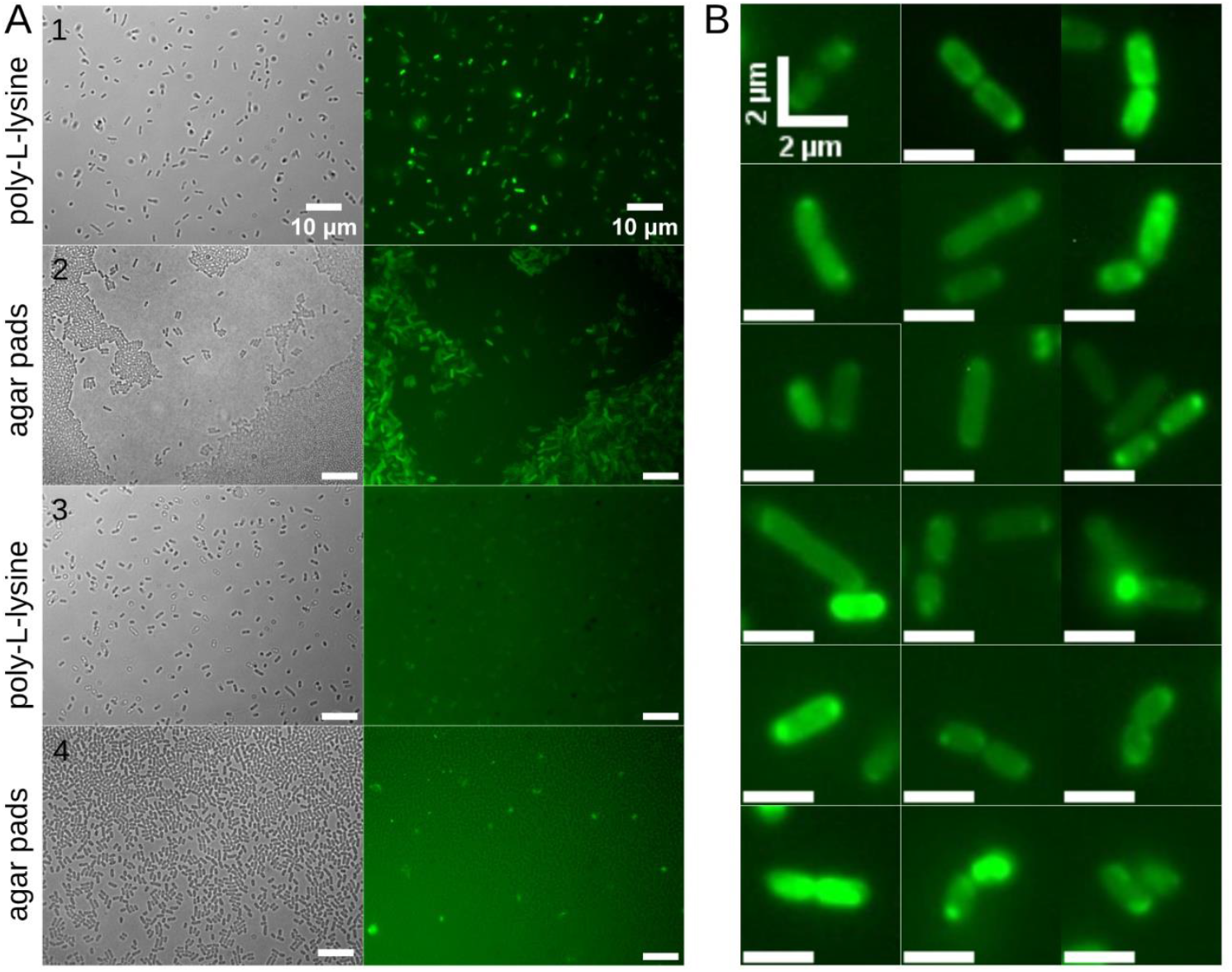
Fluorescent mapping of cardiolipin-enriched regions in *E. coli* membranes. Green fluorescence signal from 200 nM 10-nonyl acridine orange bromide (NAO) staining of cardiolipin in *Escherichia coli* cells. (A1) Standard culture, immobilized with poly-L-lysine. (A2) Standard culture immobilized on agar pads. (A3) Chemically modified culture grown in the presence of 0.6 M D-mannitol, immobilized with poly-L-lysine. (A4) Chemically modified culture, immobilized on agar pads. Subcellular localization of cardiolipin-enriched membrane regions was observed exclusively in standard cultures. NAO staining revealed distinct polar membrane microdomains associated with cardiolipin (B).

### Mass spectrometry to analyse bacterial lipid profiles

Mass spectrometry analysis was conducted to compare the lipid profiles of *Escherichia coli* lipid extracts from *ΔclsABC* triple mutant lacking cardiolipin synthase genes and *E. coli* grown under standard conditions and with the addition of 0.6 M D-mannitol. The resulting heatmap (Fig. 5A) illustrates distinct variations in cardiolipin composition across genetic and chemical modifications in relation to standard *E. coli*. As expected, the *ΔclsABC* triple mutant does not produce cardiolipins in detectable or quantifiable amounts. Interestingly, the supplementation of D-mannitol affected the profile of cardiolipin in *E. coli* both qualitatively and quantitatively. In total, lower quantity of CLs were extracted from the mannitol-supplemented bacteria, as observed by a reduction to approximately 50% of the total peak area of all identified CL. Out of 44 cardiolipins of different acyl chain lengths and desaturation identified in extracts, 16 differed significantly between the treatment conditions. The ones exhibiting the largest magnitude of change in their relative levels between experimental groups are shown in Fig. 5B. The main cardiolipin of the *E. coli* membranes was CL66:2, and its content did not change significantly between the cultures (Fig. 5A), but there were many for which levels were reduced in the mannitol-supplemented bacteria (Fig. 5B). On the other hand, it seems that longer acyl chain fatty acids (18+) and odd-numbered fatty acids were recruited into cardiolipins in the mannitol treated *E. coli*. The fatty acids in cardiolipins which were significantly elevated in the mannitol-supplemented cultures were also more unsaturated. (Fig. 5B III, IV, V, IV).

**Fig 5.**
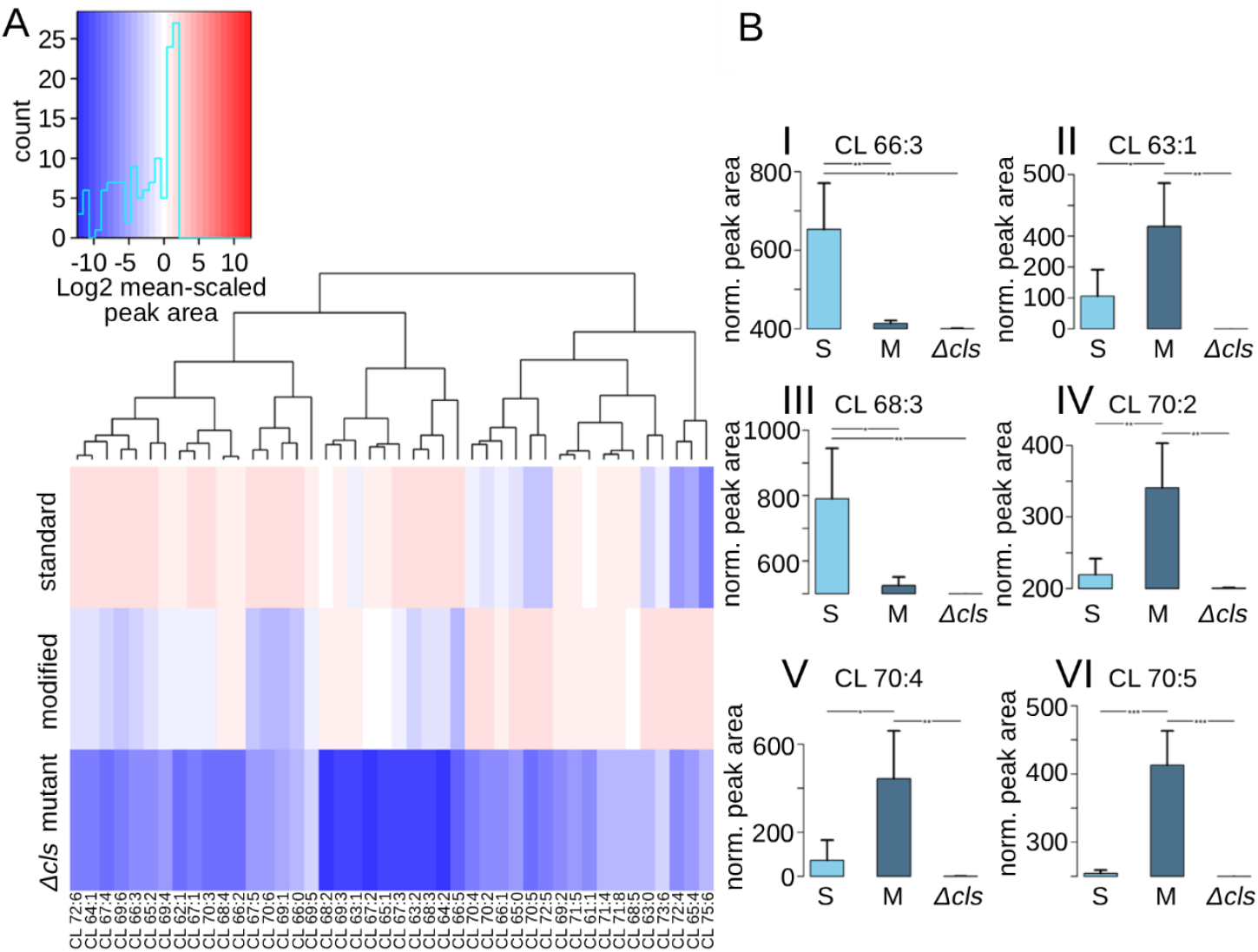
Comparison of cardiolipin species across experimental strains (A) heatmap of variations in the cardiolipin species across three experimental conditions. The figure represents normalized, scaled to experiment mean and log2-transformed peak areas. **(B) comparison of selected cardiolipin species across three experimental conditions**. Bars represent normalized mean peak area of three biological replicates with each in three technical replicates. Error whiskers represent corresponding standard deviation. ANOVA test with the TukeyHSD post-hoc analysis was used for multiple comparisons of group means and to identify significant group differences (*,**,*** represent p-value less than 0.05, 0.01 and 0.001, respectively). Labels: *Δ*cls, a *ΔclsABC* triple mutant lacking cardiolipin synthase genes; S - standard; M - modified, strain with 0.6 M D-mannitol.

### Acyl chain composition and saturation degrees of bacterial lipid extracts

To investigate the degree of fatty acid saturation, gas chromatography (GC) was performed on fatty acid methyl esters (FAMEs) obtained from total lipid extracts of *Escherichia coli* cultured under standard and mannitol conditions (fig. 6). For quantification, the internal standard used was triheptadecanoin (TG 17:0), enabling accurate comparison of lipid content between samples. The relative abundance of each detected fatty acid species is indicated next to the corresponding peaks, in both absolute values and as a percentage of total fatty acids. Notable differences were observed between the two profiles, particularly in the relative amounts of monounsaturated fatty acids. Mannitol treatment induced a substantial shift across all identified fatty acids. Approximate two-fold shift occurred in myristic acid (14:0), pentadecanoic acid (15:0), palmitoleic acid (16:1n-7), and vaccenic acid (18:1n-7). In the standard culture sample, 16:1n-7 constituted 64.90% of the total fatty acids, compared to 30.25% in the mannitol-treated culture. Similarly, 18:1n-7 decreased from 2.39% to 1.10%. Conversely, the relative abundance of saturated fatty acids such as palmitic acid (16:0) and stearic acid (18:0) showed a corresponding increase to 46,77% and 2,79% respectively.

**Fig 6.**
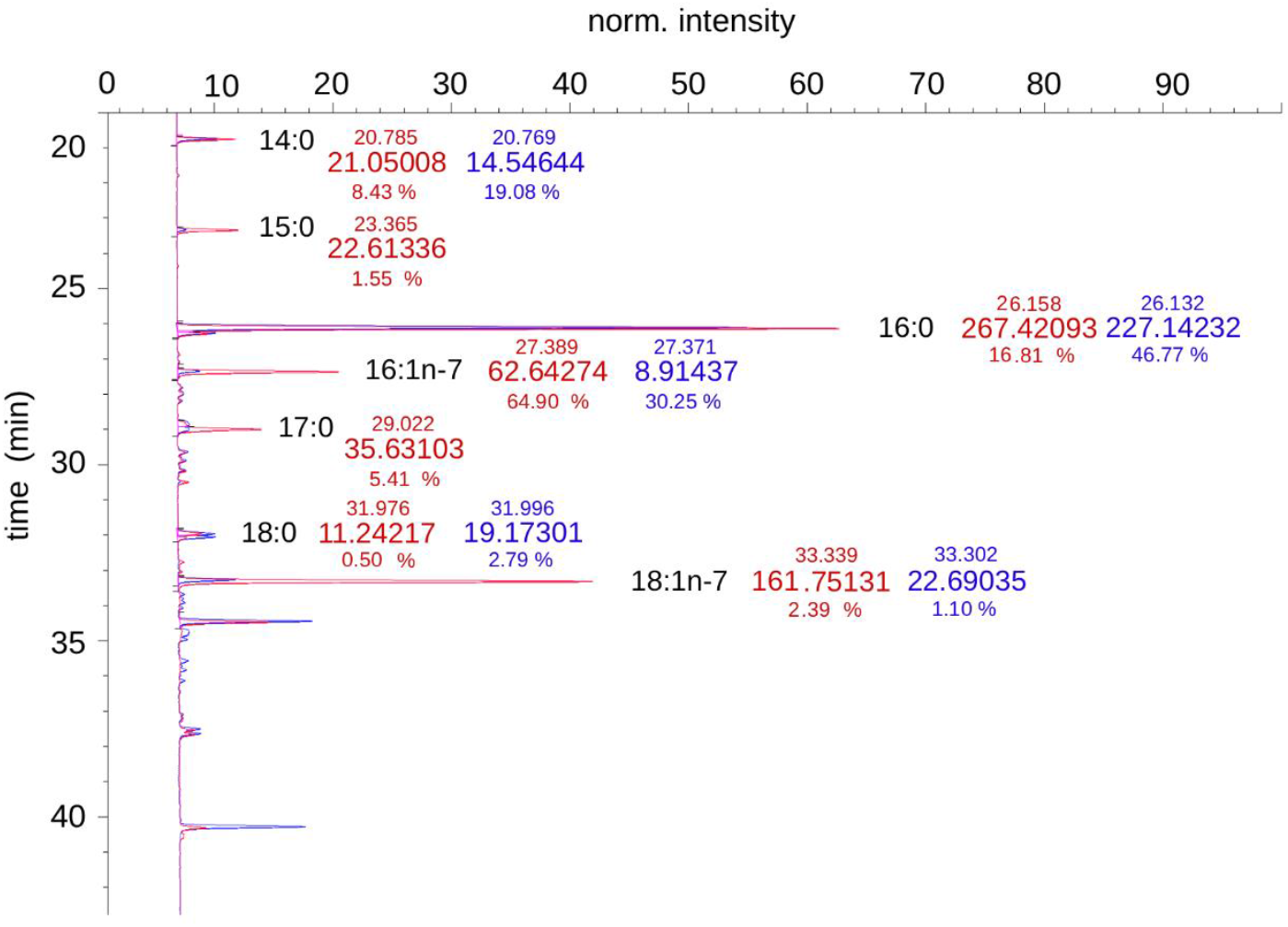
GC profiles of fatty acid methyl esters (FAMEs) derived from total lipid extracts of *Escherichia coli* cultured under standard conditions (red) and in the presence of 0.6 M D-mannitol (blue). The relative abundance (percentage of total fatty acids) for each major peak is indicated. Quantification was performed using triheptadecanoin (TG 17:0) as an internal standard. Mannitol present resulted in a marked decrease in monounsaturated fatty acids, particularly 16:1n-7 and 18:1n-7, and an increase in saturated species such as 16:0 and 18:0.

## Discussion

### APDT on *Escherichia coli* with impaired cardiolipin production

Cardiolipin is a central determinant of bacterial membrane organization, dynamics, and stress adaptation, supporting the activity and spatial organization of numerous membrane-associated proteins complexes, including SPHS proteins (QmcA, YqiK, HflK, HflC)^32^, ammonium channels AmtB,^33^ Sec translocase (SecA-SecYEG),^34^ formate dehydrogenase-N,^35^ and transporters such as LeuT.^28,36^ Although its contribution to membrane physiology has been increasingly studied, the role of cardiolipin in bacterial responses to antimicrobial photodynamic therapy remains largely unexplored.^26^ In this research, we identify cardiolipin as a critical modulator of bacterial susceptibility to aPDT and demonstrate that both genetic depletion and environmentally induced remodeling of cardiolipin profoundly sensitize *Escherichia coli* to photodynamic inactivation.

Both the *ΔclsABC E. coli* mutant and culture with modified membrane composition induced by mannitol supplementation showed significantly increased susceptibility to aPDT indicating impaired defences against photochemical reactions and oxidative stress.

### The electrokinetic properties of bacterial cell surface

Analysis of electrokinetic membrane properties, critical for maintaining membrane architecture, regulating transmembrane potential, and sustaining metabolic activity, revealed that mannitol supplementation changes the surface properties.^37^ Zeta potential measurements showed that supplementation of D-mannitol modulates the electrokinetic profile of bacterial membranes, suggesting altered interactions with the cationic photosensitizer independent of electrostatic forces. These effects are likely due to reorganization of the lipid matrix and loss of CL-rich microdomains essential for redox homeostasis and respiratory chain function. The observed shift in zeta potential value suggests a reduction in the overall negative charge on the bacterial surface, due to alterations in the composition or distribution of surface-exposed molecules, particularly the highly anionic lipid cardiolipin in mannitol-supplemented *E. coli* cultures.^38^ Many antimicrobial agents, including photosensitizers such as methylene blue, carry a positive charge. Therefore, a reduction in the negative surface charge of bacterial cells would be expected to decrease electrostatic interactions with methylene blue, potentially reducing its efficacy. Surprisingly, cultures supplemented with D-mannitol showed increased sensitivity to methylene blue, suggesting that factors beyond surface charge, such as changes in membrane composition or organization, may enhance photosensitizer activity. Importantly, supplement of D-mannitol did not significantly impact the culture conductivity. This counterintuitive result indicates that factors beyond surface charge, including the structural organization of the lipid matrix and the integrity of cardiolipin-rich microdomains, dominate the cellular response to aPDT. The increased sensitivity of *ΔclsABC* mutant to methylene blue alone (no light exposure), supports a role for cardiolipin in maintaining respiratory chain function and cellular redox balance. One possibility is that methylene blue might act as a transient, non-specific acceptor of electrons from the respiratory chain. The observed effects with changes in cardiolipin composition highlight the importance of cardiolipin in maintaining the integrity of the respiratory chain and the ability to preserve the optimal energy state of the bacterial cell.^28,39^

### Photodynamic response of biomimetic bacterial membrane model

Among the various biomimetic models available for studying biological membranes, giant unilamellar vesicles (GUVs) were particularly advantageous to investigate changes in photochemical responses of bacterial lipids under controlled conditions.^40,41^ Their utility in investigating the mechanisms of photodynamic therapy lies in their ability to replicate the lipid environment of cell membranes, enabling the assessment of photosensitizer, membrane interactions, oxidative damage, and membrane stability during light-induced stress.^42,43^ The GUV assembled from standard culture, characterized by a higher cardiolipin content and increased total acyl chain unsaturation, were found to be more susceptible to photoinactivation compared to those derived from mannitol-supplemented cultures. This observation is consistent with the well-established vulnerability of cardiolipin and unsaturated fatty acyl chains to reactive oxygen species.^39,44–46^ The increased level of cardiolipin in the extract from standard culture, comparing to mannitol-supplemented culture extract, may enhance ROS interactions, leading to oxidative damage that manifests as rapid contrast decay and structural destabilization of the vesicles.^47^ Furthermore, unsaturated fatty acyl chains are particularly vulnerable to ROS-induced peroxidation due to the presence of double bonds, which serve as primary targets for oxidative attack. This increased susceptibility contributes to membrane destabilization and may further amplify the oxidation effect.^48^

### Living cells vs. membrane model

The GUV results contrasts with the response observed in living cells, underscoring the importance of cardiolipin within the full cellular context. It reveals how living systems leverage specific lipid components to mediate stress responses and resist photodynamic therapy. In living cells, cardiolipin appears to influence aPDT susceptibility, potentially by stabilizing critical membrane-associated proteins or structural components during exposure to ROS.^25,26,39^ Such stabilization may involve disruption of membrane curvature, disruption of membrane integrity which is associated with altered bacterial responses to aPDT. A higher cardiolipin content in standard culture may enhance membrane resilience by supporting proteins involved in energy metabolism, transport, and repair that are essential under oxidative stress conditions.^29,39^ Conversely, the reduced cardiolipin content in mannitol-supplemented cells may impair membrane organization and folding make bacteria more vulnerable to ROS damage due to lack of cardiolipin-rich microdomains and the support of ROS defence systems.^29,49^

### Cardiolipin visualisation by fluorescence microscopy

Integration of the cardiolipin-rich microdomains visualization and localization data with findings on aPDT susceptibility in lipid extracts and live *E. coli* cells gives valuable insight into how cardiolipin distribution and relative abundance influence the cell membrane properties. In standard culture, CL is prominently localized at the poles of the inner membrane, forming distinct cardiolipin-rich microdomains.^31,50,51^ These specialized regions in standard-culture cells could enhance their resilience to ROS generated during aPDT by creating microenvironments that support cellular ROS defence mechanisms. Such structural organization could explain why standard-culture cells exhibit reduced susceptibility to aPDT, despite their high cardiolipin content. In mannitol-supplemented cultures, where cardiolipin level is significantly reduced and its structure is modified, fluorescence microscopy reveals no CL signal even on poles regions. This suggests a redistribution or depletion of CL from key structural regions of the membrane, potentially altering membrane fluidity and compromising its ability to support membrane proteins critical for stress defence thus diminishing support for ROS-scavenging enzymes or transport systems, leading to an increased susceptibility to oxidative damage during aPDT.^28,29^ This reduction and remodelling of cardiolipin at the cell poles could explain why mannitol-supplemented cells are more sensitive to photoinactivation in live-cell experiments, as the membrane may lack the stabilizing influence of cardiolipin-rich microdomains.

### Mass spectrometry to analyse bacterial lipid profiles & acyl chain composition

Mass spectrometry analysis confirmed that both genetic and environmental modifications significantly alter CL abundance and composition in *Escherichia coli*. The observed substantial reduction in total cardiolipin (CL) abundance in the mannitol-supplemented culture aligns with established mechanisms of membrane adaptation to stress. Furthermore, the lack of detectable or quantifiable amounts of CL in the *ΔclsABC* triple mutant is consistent with the essential role of these three genes (*clsA, clsB, clsC*) in the primary CL synthesis pathway in *Escherichia coli*, thereby validating the genetic control used in this study.^30,31^ Under standard conditions, bacteria exhibited a diverse cardiolipin profile, dominated by CL66:2 species. Interestingly, the addition of 0.6 M D-mannitol to the culture medium resulted in a significant (∼50%) decrease in total cardiolipin content, with 16 of 44 species showing significant changes. The content of several shorter-chain and less unsaturated cardiolipins was reduced, whereas species incorporating longer acyl chains (≥ C18) and odd-numbered fatty acids were relatively enriched. Moreover, cardiolipins elevated under mannitol conditions showed a higher degree of desaturation. Comprehensive determination of the fatty acid saturation of the extracted lipids by GC showed an overall reduction in monounsaturated fatty acids, in particular palmitoleic acid (16:1n-7) and vaccenic acid (18:1n-7).

Integration of lipidomic, microscopy, and functional data reveals that not only cardiolipin abundance but also its spatial distribution is critical for bacterial resistance to aPDT. Findings indicate that cardiolipin, as a previously unrecognized determinant of bacterial susceptibility to antimicrobial photodynamic therapy, plays here a key role. CL biosynthesis and remodelling are strictly linked to key structural and functional organization of bacterial membrane that enhances bacterial resilience in living systems. These insights provide a new framework for understanding membrane-mediated responses to aPDT and suggest that targeting cardiolipin may represent a promising strategy to potentiate aPDT against bacterial pathogens.

## Conclusions

Cardiolipin appears to function as a multi-dimensional mediator in aPDT sensitivity. Its primary role lies in maintaining structural resilience and metabolic support, impaired in both the CL-deficient *ΔclsABC* mutant and CL-reduced mannitol culture, and observed in their increased susceptibility to aPDT. The distribution of CL across cell membrane is strictly defined by forming distinct cardiolipin-rich microdomains, localized at the poles of the membrane, where it stabilizes key protein complexes involved in transport systems, energy metabolism and cell division. CL content modulates the electrokinetic properties of the bacterial membrane. The increased sensitivity to MB occurs despite a reduction in negative surface charge, indicating that structural and functional alterations of the membrane outweigh electrostatic forces. The higher sensitivity to MB alone in absence or impartment of CL indicate its importance in supporting redox homeostasis and respiratory chain function under stress. Last but not least, while CL stabilizes the cell, its chemical nature makes it a prime target for the damaging effects of aPDT. This inverse susceptibility trend between GUVs and live-cell assays highlights that CL is not only as a passive ROS target but as an active scaffold stabilizing protein-lipid complexes, modulating membrane curvature, and anchoring stress-responsive pathways *in vivo*. Together, these findings emphasise the importance of cardiolipin in aPDT sensitivity. and suggest that modulating membrane lipid composition may potentiate the efficacy of aPDT against Gram-negative pathogens.

## Methodology

### Bacterial strains and growth conditions

*Escherichia coli* PCM 172, obtained from the Polish Collection of Microorganisms, served as the model organism and reference strain for all experiments. Cultures were grown under optimal conditions at 37°C with agitation at 180 rpm in nutrient broth containing 10 g/L Lab-Lemco powder, 10 g/L peptone, and 5 g/L NaCl, adjusted to a pH of 7.5 ± 0.2. To induce alterations in membrane lipid composition, the medium was supplemented with 600 mM D-mannitol (commercially sourced). The *ΔclsABC* mutant, characterized by deletions in the cls genes, was obtained from Carranza et al. ^30^

### Antimicrobial photodynamic therapy evaluation

APDT was evaluated on planktonic bacterial cultures with or without 0.6 M D-mannitol in a 96-well plate format (250 μL per well), following the methodology described by Piksa et al. 2023 ^6^ Regardless of the growth conditions, bacterial cultures were standardized to an optical density (OD) of 0.001. Three experimental groups were prepared: (1) a growth control (C) containing a bacterial suspension without photosensitizer (no PS, non-irradiated), MB group a photosensitizer-only control to assess the effect of the photosensitizer in the dark (with PS, non-irradiated), and aPDT group containing methylene blue and exposed to OLED irradiation (with PS, irradiated). The final concentration of methylene blue in the bacterial suspension was 15 μM.

The aPDT was conducted at 37 °C with continuous illumination of the aPDT group for 3 hours. The OLED light source was positioned approximately 4 mm above the plate, delivering an energy dose of 54 J/cm^2^ at an irradiance of 5 mW/cm^2^. Each experimental condition used a single bacterial suspension (representing one biological replicate) divided into three wells, providing three technical replicates (n = 3). In total, three biological replicates were performed, yielding nine technical replicates per condition (n = 9). Following aPDT, the plate was centrifuged at 4750 × g for 5 minutes, and the supernatant was replaced with fresh medium.

Bacterial growth measurements were carried out as described by Lian et al.^47^using a CLARIOstar® Microplate Reader (BMG LABTECH). The plate was incubated at 37 °C in the dark with continuous double orbital shaking at 300 rpm for at least 20 hours. Absorbance at 450 nm was recorded every 15 minutes to generate detailed growth curves. Inhibition by methylene blue or aPDT was indicated by a slower increase in absorbance over time and a lower final absorbance. Results were compared using the high-throughput Start-Growth-Time (SGT) method.

### Zeta-potential of bacterial cell surface

Bacterial culture under standard condition with or without D-mannitol grew to obtain OD=1.0 and centrifuged 3500 rpm 20min 20°C. Supernatant was removed and bacterial pellet was washed 3 timed on PBS with centrifuging each time (4500 rpm 20min 20°C). After last wash and centrifuging bacteria in 1 mL of PBS, bacterial suspension was placed in special cuvette and measurements were conducted. Each measurement was repeated 3 times and 3 biological replications were prepared (n=9).

### Giant unilamelar vesicles from bacterial lipid extract

Giant unilamellar vesicles (GUVs) were prepared separately from *Escherichia coli* standard culture and D-mannitol-supplemented culture according to a modified method from Weinberger et al. (Gel-Assisted Formation of Giant Unilamellar Vesicles). Firstly, the layer of PVA was prepared by placing 150 μL of poly(vinyl alcohol) (PVA) (5%; RT) on a solid surface and keeping it in the oven (70°C) for 60 min and cooling down for 10 minutes (RT). 10 μL of lipid solution in chloroform (1 mg/mL) was placed on PVA layer and dried in a desiccator under vacuum pressure for min. 60 min. Next, 150 μL of 100 mM sucrose solution (RT) was added and to induce GUVs growth and left for min. 50 min in RT. After growing time GUVs were collected carefully and slowly with tips (cut 0.5 cm of end) or syringe (no needle) and put in Eppendorf tube (around 150 μL). 1350 μL of 90 mM glucose solution (RT) was added slowly and left for 30 min to let GUVs sediment. 55 μL of GUVs from the bottom were placed inside the observation cell on the cover slip and 55 μL of 30 mM methylene blue (MB) solution in 90 mM glucose was added. The samples prepared as described above were subjected to aPDT conditions directly under the microscope with Zeiss Axio-observer 7, the inverted microscope in phase contrast mode equipped with an Orca-Flash 4.0 LT camera (Hamamatsu) and Colibri 7 fluorescent LED illumination source (631 nm) was used.

### Cardiolipin visualisation by fluorescence microscopy

Bacteria were incubated with 10-nonyl bromide acridine orange (NAO, Sigma-Aldrich) (200 nM) NAO. Two immobilisation methods were used. The first was based on 1% agar pads and the second on 0.01% poly-L-lysine. In both cases, 10 μL of bacterial culture was transferred directly on immobilization area. The fluorescent dye was excited using a Zeiss HBO 100 Fluorescence Microscope Lamp. Green fluorescence (with excitation at 495 nm and emission at 525 nm) from NAO was detected by using a standard GFP filter unit (Ex. 470/40–Em. 525/50). Standard fluorescence microscopy observations were carried out using Zeiss Axio Imager upright fluorescence microscope (Zeiss EC Plan-Neofluar 100/1.30 Oil) equipped with a 12 bits B&W camera (AxioCam MRm). The image analysis was performed with the use of ImageJ v.1.54f.

### Lipid extraction

Bacterial pellets were extracted using 1000 μl of cold (-20 °C) mixture of methyl-*tert*-butyl-ether : methanol (3:1, v/v) with the addition of internal standards (0.1 μg/ml deuterated phosphatidylcholine (PC36:0-D_70_) and 0.1 μg/ml deuterated arachidonic acid (AA-D_5_)). The samples were then sonicated using a cooled (4 °C) ultrasonic bath for 10 min. Then, 500 μl mixture of water : methanol (3:1, v/v) was added to each of the samples, which led to the formation of two liquid phases – polar and nonpolar. The upper lipid phase was collected into a new tube, dried with a speedvac and stored at -20 °C, prior to the lipidomic profiling.

### Lipidomic analysis

The nonpolar phase lipids were analyzed using a Waters Acquity UPLC system coupled with a Xevo G2 QTof mass spectrometer. The LC conditions were column ACQUITY UPLC BEH Shield RP18 (2.1 x 100 mm, 1.7 μm, Waters); column temp. 60 °C; mobile phase A acetonitrile/water (60:40, v/v) with 10 mM ammonium formate and 0.1% formic acid; mobile phase B isopropanol/acetonitrile (90:10, v/v) with 10 mM ammonium formate and 0.1% formic acid; injection volume 5 μl. A constant flow rate of 300 μl/min was maintained over 25.5 minutes of an analytical run. Following mobile phase gradient was used: initial 70.0% A, 3.0 min. 70% A, 6.5 min. 55% A, 12.5 min. 40% A, 19.0 min. 5% A, 21.5 min. 5% A, 21.51 min. 70% A, 25.5 min. 70%. The mass spectrometry conditions were: ESI capillary voltage 1kV (negative), source temp. 120 °C, desolvation temperature 450 °C, acquisition mode MS^E^ (low CE 10V, high CE 30V), scanning range 100-1800 m/z. Eluents were Hypergrade for LC-MS quality purchased at Merck. Raw acquired chromatograms were converted to .abf files by Reifycs Analysis Base File Converter with default settings for Waters’ MS^E^ files. For peak detection, alignment, annotation, peak area integration, the .abf files were loaded into MS-DIAL software (version 4.9.; RIKEN Institute). Additionally the search and identification was supported by use of XCMSonline tool (https://xcmsonline.scripps.edu) for which the raw data were converted to .mzXML format with the MSConvert tool (version 3.0.23051-d77d375). The output file was further compared to the *in-house* database of lipids for refined manual annotation of peaks. Mixtures of cardiolipin species (CAS 796964-05-1, 841199P, Avanti) was used to support annotation.

In order to analyse the saturation of the acyl chain after adding appropriate internal standards to the samples, a fraction of total lipids was transmethylated with boron trifluoride in methanol (14%). Tubes was carried out at 100 °C during 1H30. The derivatized fatty acid methyl esters were then extracted twice with isooctane and analyzed by gas chromatography using an HP6890 instrument equipped with a fused silica capillary BPX70 SGE column (60 mx 0.25 mm). The vector gas was hydrogen. Temperatures of the flame ionization detector and the split/splitless injector were set at 250 °C and 230 °C, respectively.

